# Detecting space-time clusters of dengue fever in Panama after adjusting for vector surveillance data

**DOI:** 10.1101/561902

**Authors:** Ari Whiteman, Michael R. Desjardins, Gilberto A. Eskildsen, Jose R. Loaiza

## Abstract

Long term surveillance of vectors and arboviruses is an integral aspect of disease prevention and control systems in countries affected by increasing risk. Yet, little effort has been made to adjust space-time risk estimation by integrating disease case counts with vector surveillance data, which may result in inaccurate risk projection when several vector species are present, and little is known about their likely role in local transmission. Here, we integrate 13 years of dengue case surveillance and associated *Aedes* occurrence data across 462 localities in 63 districts to estimate the risk of infection in the Republic of Panama. Our space-time modelling approach detected the presence of five clusters, which varied by duration, relative risk, and spatial extent after incorporating vector species as covariates. Dengue prevalence (n = 49,910) was predicted by the presence of resident *Aedes aegypti* alone, while all other covariates exhibited insignificant statistical relationships with it, including the presence and absence of invasive *Aedes albopictus*. Furthermore, the *Ae. aegypti* model contained the highest number of districts with more dengue cases than would be expected given baseline population levels. This implies that arbovirus case surveillance coupled with entomological surveillance can affect cluster detection and risk estimation, improving efforts to understand outbreak dynamics at national scales.

**Author Summary:** Dengue cases have increased in tropical regions worldwide owing to climate change, urbanization, and globalization facilitating the spread of *Aedes* mosquito vectors. National surveillance programs monitor trends in dengue fever and inform the public about epidemiological scenarios where outbreak preventive actions are most needed. Yet, most estimations of dengue risk so far derive only from disease case data, ignoring *Aedes* occurrence as a key aspect of dengue transmission dynamic. Here we illustrate how incorporating vector presence and absence as a model covariate can considerably alter the characteristics of space-time cluster estimations of dengue cases. We further show that *Ae. aegypti* has likely been a greater driver of dengue infection in high risk districts of Panama than *Ae. albopictus*, and provide a discussion of possible public health implications of both spatial and non-spatial model outcomes.

## Introduction

Dengue fever, a disease transmitted to humans by *Aedes* mosquitoes, is endemic to 128 countries, with 3.9 billion people considered at-risk [1]. Dengue fever cases have increased dramatically worldwide throughout the previous several decades [2], likely a result of climate change [3], urbanization [4], globalization [5], and the spread of the invasive *Aedes albopictus* [6]. As a result of both recent and historical risk, many countries employ national surveillance programs to monitor trends in dengue fever and inform local health authorities to the places and times where preventative practices are most required. However, despite the commonality of these programs and unforeseen cost of cutting them [7], surveillance budgets are often limited [8,9], restricting the scope and quality of the work. This is concerning in developing regions such as Central America, where the burden of disease is high [1] and per capita public health expenditure is among the lowest of any region of the world [10].

Surveillance of both viruses and vectors is an essential component of integrated disease management programs that can be used to determine risk changes in space and time, thus providing the evidence for more targeted prevention and control interventions [11]. Nevertheless, with few exceptions, it is rare for surveillance programs to concurrently monitor both arbovirus cases and vector populations in the same locations and at regular intervals. Most projections of disease risk used to justify public health actions are derived purely from disease case data, ignoring vector population dynamics, which is key aspect of the vector transmission model. This is particularly concerning when more than one vector species is present, and little is known about their likely role in local transmission, which may result in inaccurate or incomplete risk projection or case clustering models.

The Republic of Panama has been monitoring dengue cases alongside vector presence through the National Department of Epidemiology (NDE) since 1988, making it one of the most long-standing and successful surveillance programs of its kind in Latin America. Of the two known dengue mosquito vectors, *Ae. aegypti* is considered resident to Latin America and Panama since the 19^th^ century, and the primary source of transmission [11] while *Ae. albopictus*, considered a secondary vector, has been spreading throughout the region ever since it got introduced in Panama in 2004 [12,13]. Widespread extirpation of *Ae. aegypti* by a superior ecological competitor like *Ae. albopictus* has occurred throughout the world in recent decades [14–16], with unknown consequences on arbovirus transmission risk. Encompassing this period of growing interspecific competition among two vector species, Panama’s surveillance system is particularly unique and potentially useful to modelling dengue transmission risk while considering *Aedes* species interaction. Attaining a better understanding of dengue outbreak dynamics over time may improve the capacity of public health authorities to combat the spread of other arboviruses, such as Zika Virus and Chikungunya Virus.

Our overall aim is to examine the influence that concurrent dengue case surveillance and *Aedes* species monitoring can have on cluster detection and relative risk estimation. In so doing, we describe the results of 13 years of dengue fever and *Aedes* surveillance data, including two competing vector species plus virus data originating from long-term cooperatively organized surveillance programs. We further assess whether dengue prevalence can be attributed to district socioeconomic attributes. We believe this is the first effort to adjust for vector presence and absence in a disease cluster detection model, which we hope sheds light on the characteristics of space-time clusters and relative risk estimation of dengue after *Aedes* species are used as model covariates.

## Methods

### Dengue Data

We utilized dengue prevalence data collected by the National Department of Epidemiology (NDE), housed within the Panamanian Ministry of Health (MINSA). Systematic national surveillance of dengue cases in Panama have been continuous since 1988. Suspected cases are defined by a patient with a fever and one or more of the following symptoms: headache, retro orbital pain, myalgia, exanthema, rash, vomiting, malaise, leukopenia, and jaundice. A confirmed case is defined as a suspected case with a positive dengue test, conducted using either viral isolation, reverse transcription polymerase chain reaction (RT-PCR), IgM enzyme-linked immunosorbent assay platform (ELISA), or secondary IgG ELISA. RT-PCR was established as the original standard by the National Reference Laboratory at the Gorgas Memorial Institutes for Health Studies (ICGES) in 2003. Yet since 2009, MINSA established national decentralization of serological confirmation of dengue using ELISA tests, which has improved efficiency by allowing district health officials to confirm cases without needing to send samples to a single central facility in Panama City. Data is recorded at the *Corregimiento,* or neighborhood, scale as the number of confirmed cases in a given year at a given location. This is the lowest scale of data granularity available, and thus, we do not have patient-level detail nor temporal detail at smaller units than year.

### Vector Data

We utilized vector data from the Vector Control Department (VCD) at MINSA. Systematic entomological surveillance has occurred in Panama since 2000 in order to establish *Aedes* infestation rates, and thus, areas of potential dengue transmission risk. Surveys of both *Ae. aegypti* and *Ae. albopictus* are performed annually at the *Corregimiento*-scale and consist of solely larval surveillance. Each year, a random block of houses is chosen and all houses in the block are searched for containers holding *Aedes* larvae. The larvae are collected and allowed to mature to the fourth instar, at which point they are taxonomically identified to species based on morphological keys [17]. The number of houses positive for *Ae. aegypti*, *Ae. albopictus* or both are recorded in the raw datasets. However, because we cannot confirm the number of houses in each block, we have transformed the data into a presence-absence format in each *Corregimiento* rather than analyzing the number of positive houses.

### Data Analysis

We conducted our analyses on dengue and vector data from 2005-2017, encompassing the period in Panama when both *Ae. aegypti* and *Ae. albopictus* have been interacting. Overall, data was collapsed from the original *Corregimiento* scale to the district scale. This is due to unreliable human population estimates at scales smaller than the district. Population levels were required to compute prevalence rate (x1000; PR), which was used as the dependent variable in the statistical analysis, rather than pure number of cases, which does not consider the total number of potential virus hosts. Human population data was gathered from the National Institute of Statistics and Census (INEC), which conducts a national census every 10 years. We also gathered three socioeconomic metrics from INEC to use as covariates: percentage of households with dirt floors, percentage of households without clean water, and percentage of households without sanitary services. These covariates were chosen due to their relationship to standing water, which may act as potential *Aedes* breeding habitat. Because the national census is only conducted every ten years, we used the population levels from 2010 to calculate PR for data from 2005-2017. While this is not ideal, and incurs inherent error in the year to year accuracy of the PR estimate, there is no more frequent population estimate available. This is an unfortunately common situation, especially in Central America, where no country conducts national population assessments more frequently than every 10 years. The three socioeconomic variables are at their 2010 levels as well, sourced from the same census as population.

We conducted two sets of analyses, non-spatial and spatial. The purpose of the spatial analyses was exploratory, assessing the relationship between vector and virus in space and time. This was conducted first, to establish a baseline understanding of how the addition of vector surveillance data affects the estimation of the size and relative risk of case clusters. We followed the spatial modeling with non-spatial statistical modelling, which served to test the hypotheses established by the spatial models. Thus, the non-spatial models essentially serve to identify the significant covariates that can be adjusted for in the spatial model For the spatial analyses, we utilize discrete Poisson space-time modelling STSS [18], which systematically moves cylindrical search windows across the geographic and temporal space to detect space-time clusters. Essentially, STSS determines if the observed disease cases in a particular region and time period exceed the expected cases under baseline conditions. In vector-borne disease research, STSS have been used to examine outbreaks of dengue [19–21], chikungunya [22], malaria [23,24], Chagas [25], and West Nile [26,27], for example. STSS have also been used to examine the co-circulation of dengue and chikungunya in Colombia [28].

The cylinders are centered on the centroids of the Panamanian districts while the base of a cylinder is defined as the spatial scan, and the height of a cylinder represents the temporal scan. The number of observed and expected dengue cases are computed for each cylinder. Conceptually, a vast number of cylinders of various space-time dimensions are generated until an upper bound is reached, while each cylinder is a potential cluster. For this study, the maximum spatial scan was set to 25% of the total population in Panama, while the maximum temporal scan was set to 4 years. A Poisson-based likelihood ratio is calculated for each cylinder, which is proportional to (*n*/*μ*)^*n*^[(*N* − *n*)/(*N* − *μ*)]^*N* − *n*^ [29]. For the parameters, μ is the expected number of dengue cases in a cylinder, and n is the total observed dengue cases in the cylinder. The expected number of dengue cases is computed by multiplying the fraction of population that lives within the cylinder (*p)* by the total number of cases in Panama (*C)* divided by the total population (*P)*, that is: *E*[*c*] *= p*C/P* - The cylinder with the highest likelihood ratio is the most likely space-time cluster. To evaluate the statistical significance of the candidate space-time clusters, 999 Monte Carlo simulations are performed under the null hypothesis that there are no significant clusters. Subsequently, we report secondary space-time clusters with a p-value less than 0.05.

For this study, we ran four STSS models: (1) dengue cases only; (2) dengue cases controlled for the presence and absence of *Ae. aegypti* and/or *Ae. albopictus* (i.e. absence of both species, *Ae. aegypti* presence, *A. albopictus* presence, and presence of both species); (3) dengue cases controlled for *Ae. aegypti* presence/absence only; and (4) dengue cases controlled for *Ae. albopictus* presence/absence only. For the covariate adjusted models, the expected number of dengue cases is defined the same way for the non-adjusted model, but includes covariate category *i*. That is: 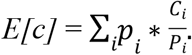 In other words, the adjusted STSS searchers for clusters “above and beyond that which is expected due to these covariates” (47). For each model, we also report the relative risk of prevalence in each district that belongs to a space-time cluster, which is defined as (*c*/*e*)/[(*C* − *c*)/(*C* − *e*)], where c is the total observed dengue cases in a particular district; *e* is the expected cases in a district; and *C* is the total observed dengue cases in the country of Panama. Clusters with a relative risk > 1 indicates that there were more observed dengue cases than expected under baseline conditions. We created all maps in ArcGIS [30].

In the non-spatial analyses we used generalized linear models (GLM; Mccullagh & Nelder, 1972) with a log linkage to determine if dengue PR could be predicted by the presence of *Ae. albopictus* alone, *Ae. aegypti* alone, the presence of both species, and the three socioeconomic attributes of the district. In addition, we tested whether the presence of *Ae. aegypti* was negatively associated with the occurrence of *Ae. albopictus*, which has been proposed by previous studies describing a pattern of spatial displacement. GLMs are robust and capable of being applied to data without homogeneous variance or normality. They have been utilized in a variety of studies on the public health implications of *Aedes* mosquito ecology [32–34].

## Results

From 2005-2017, there were a total of 49,910 cases of dengue fever in Panama, with 2009 and 2014 being the most severe at 6,941 and 7,423 cases respectively. These two years represented 28% of the total dengue cases during the 13-year period. Additionally, at the start of the sample period, *Ae. albopictus* was only present in 1 district, yet by 2017 had been found in 53 districts. It exhibited a slightly increasing trajectory throughout time and has been present in the same number of districts as *Ae. aegypti* since 2016. Surveillance of *Ae. aegypti* indicated fluctuating presence throughout the sample period, with presence ranging from 48-57 districts (Fig 1).

**Fig 1.**
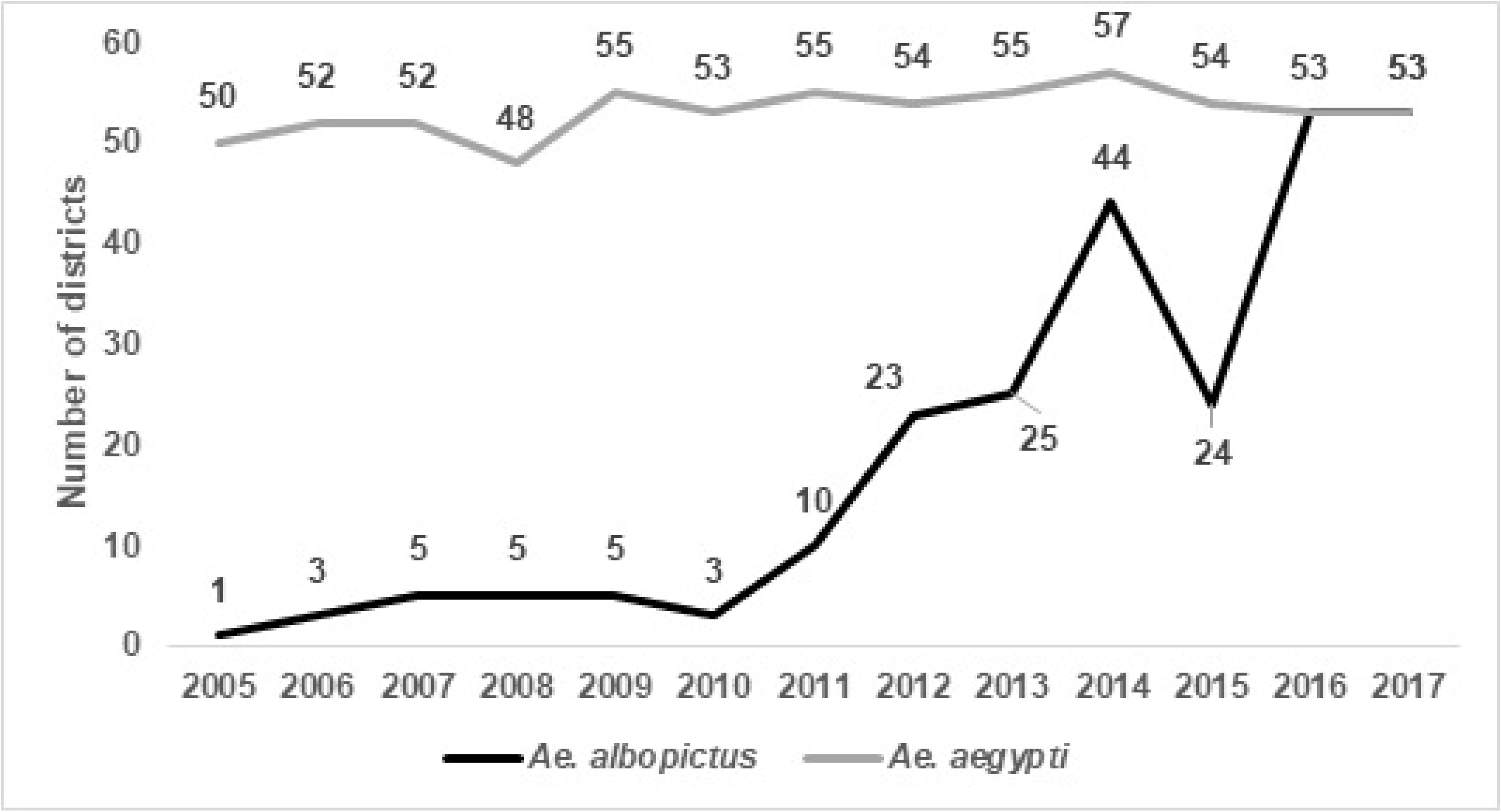
Number of districts containing each *Aedes* species from 2005-2017.

The results of our space-time modelling detected the presence of five clusters in each of the four models, varying by cluster center and duration (Figs 2-5; Table 1). Incorporating covariates into the models had considerable effects on the duration, relative risk (RR), and spatial extent of clusters (Table 2). The model adjusting for the presence of *Ae. aegypti* encompassed the greatest spatial range and highest number of districts with a RR > 1, while the model adjusting for the presence of *Ae. albopictus* encompassed the smallest spatial range and the lowest number of districts with a RR > 1. The duration of the space-time clusters is notably different when adding the vector surveillance data to the model, however, the one exception is cluster 1 for each model (most likely cluster). For example, the duration of cluster 2 was 2015-2017 for the no covariate and *Ae*. *aegypti* model; while the *Ae. albopictus* and *Aedes* (both) model reported a duration of only 1 year, which occurred six years earlier (2009).

**Fig 2.**
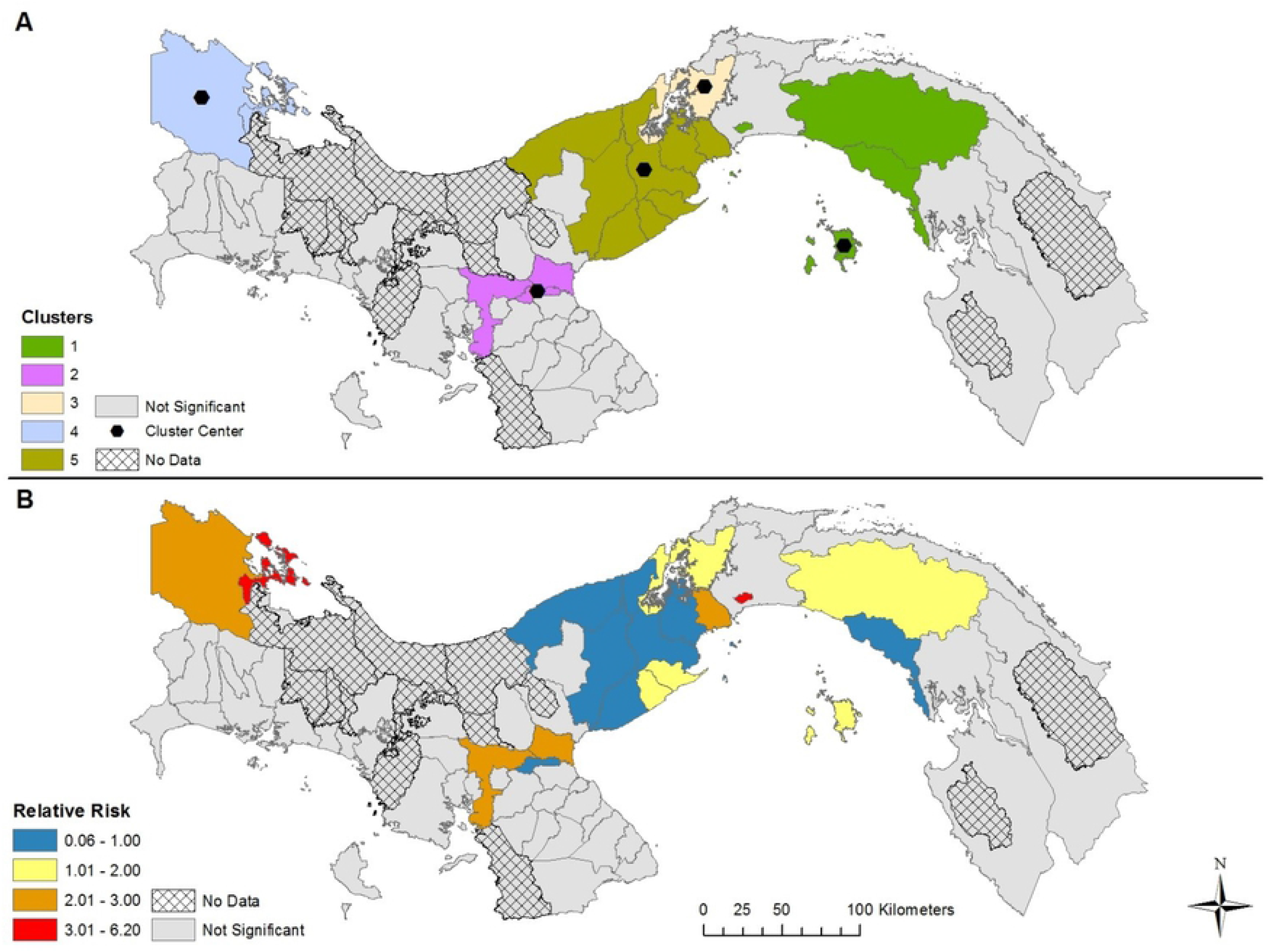
Space-time clusters of dengue fever without adjusting for *Aedes* presence and absence in Panama (A); Relative risk for districts belonging to a significant space-time cluster (B). Map created using ArcGIS [30] and data from The Panamanian Ministry of Health.

**Fig 3.**
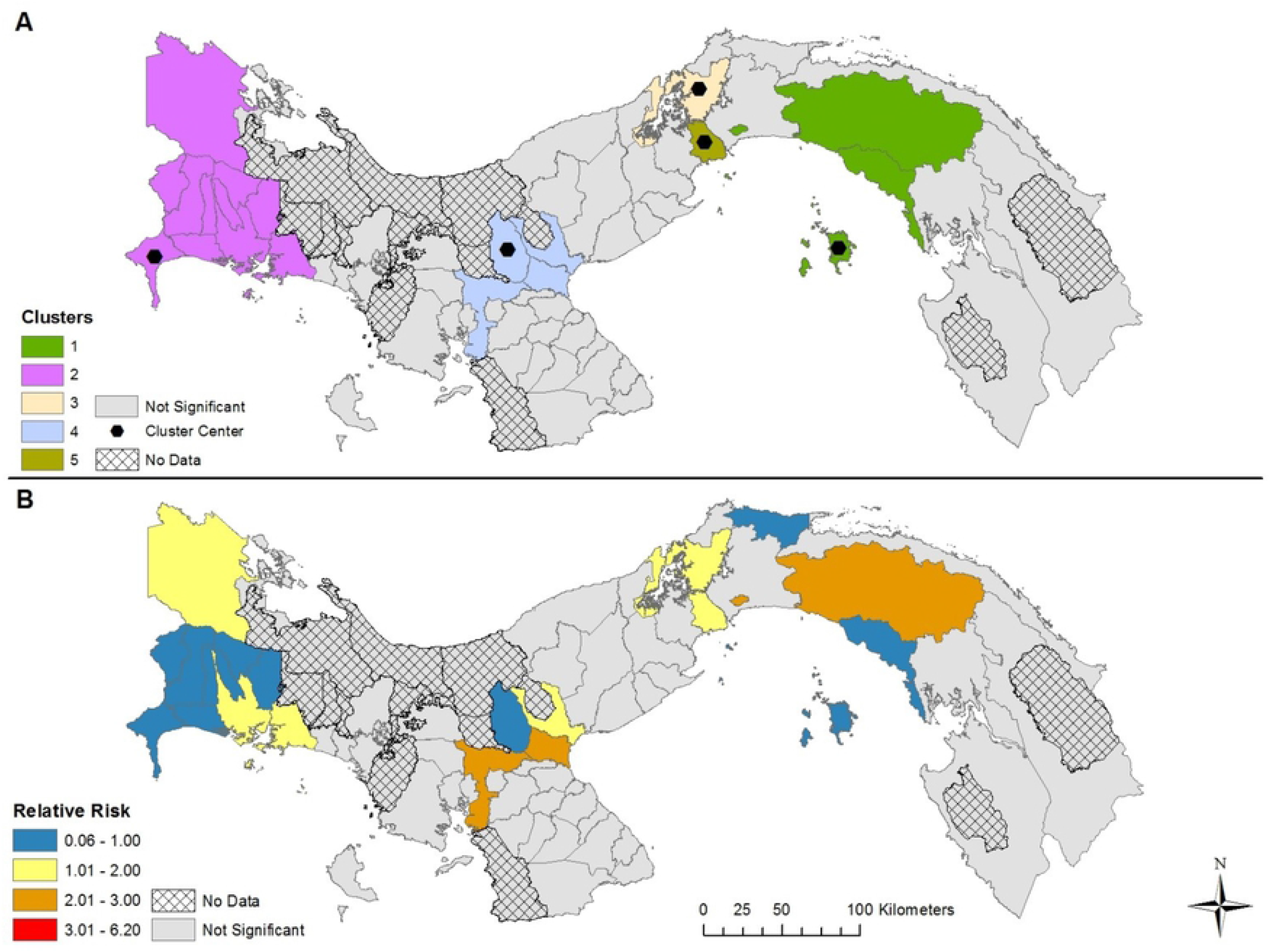
Space-time clusters of dengue that adjusts for both *Aedes* species presence and absence in Panama (A); Relative risk for districts belonging to a significant space-time cluster (B). Map created using ArcGIS [30] and data from The Panamanian Ministry of Health.

**Fig 4.**
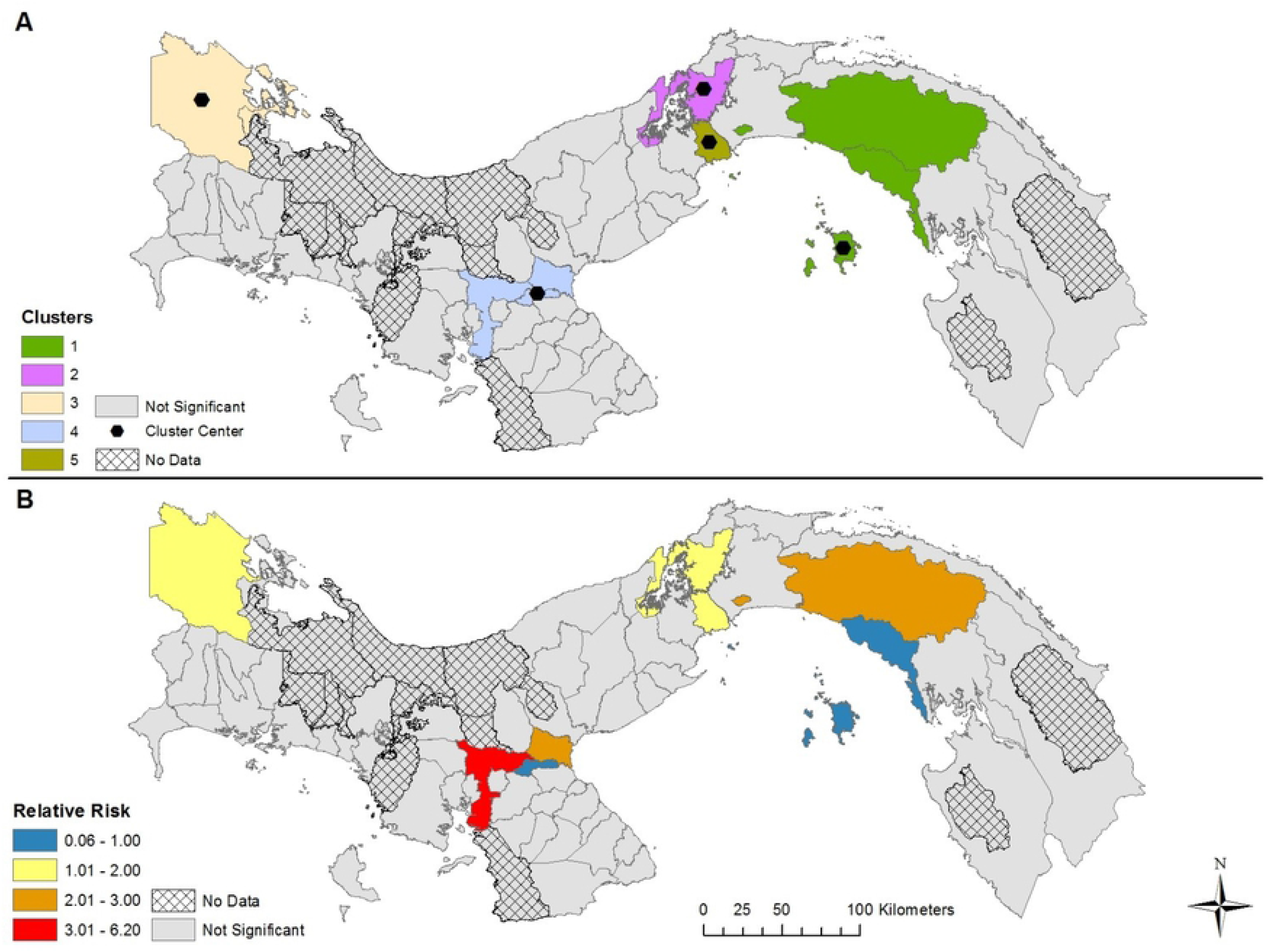
Space-time clusters of dengue fever that adjusts for *Ae. albopictus* presence and absence in Panama (A); Relative risk for districts belonging to a significant space-time cluster (B). Map created using ArcGIS [30] and data from The Panamanian Ministry of Health.

**Fig 5.**
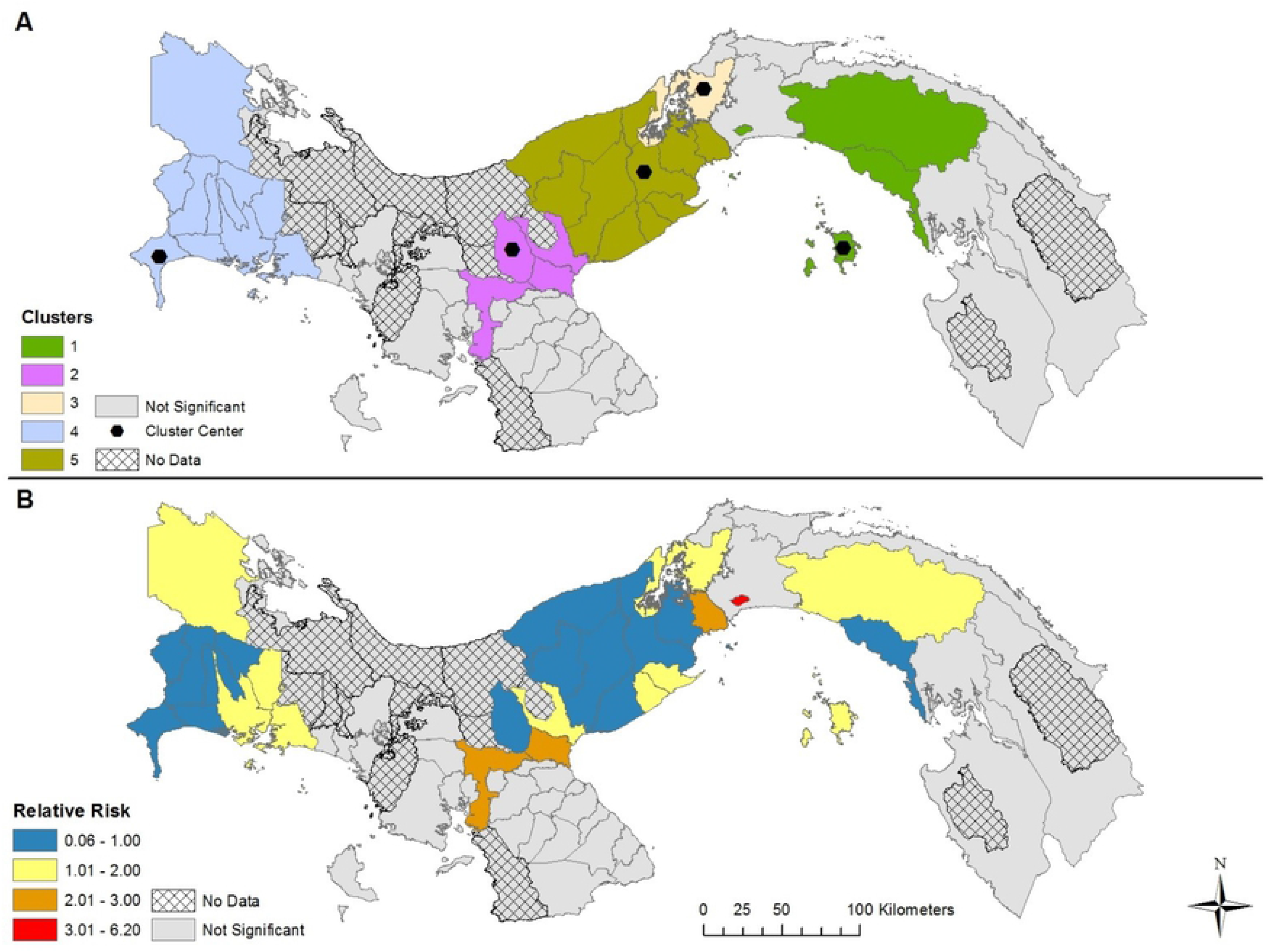
Space-time clusters of dengue fever that adjusts for *Ae. aegypti* presence and absence in Panama (A); Relative risk for districts belonging to a significant space-time cluster (B). Map created using ArcGIS [30] and data from The Panamanian Ministry of Health.

**Table 1.** Space-time dengue fever clusters.

| Cluster | Center of Cluster | Duration<br>(years) | p-value | Observed | Expected | Relative<br>Risk | Districts | Cluster<br>Population |
| --- | --- | --- | --- | --- | --- | --- | --- | --- |
| **No covariates** |  |  |  |  |  |  |  |  |
| 1 | Balboa | 2013-2015 | p<0.01 | 5,846 | 1,270.83 | 5.1 | 5 | 368,341 |
| 2 | Santa Maria | 2015-2017 | p<0.01 | 2,013 | 482.25 | 4.3 | 3 | 139,778 |
| 3 | Colon | 2009 | p<0.01 | 1,402 | 237.54 | 6 | 1 | 206,553 |
| 4 | Changuinola | 2005-2007 | p<0.01 | 1,734 | 394.85 | 4.5 | 2 | 114,445 |
| 5 | Capira | 2014 | p<0.01 | 1,914 | 721.3 | 2.7 | 9 | 627,220 |
| **Adjusting for *Aedes* presence & absence** |  |  |  |  |  |  |  |  |
| 1 | Balboa | 2013-2015 | p<0.01 | 5,846 | 1,670.20 | 3.8 | 5 | 368,341 |
| 2 | Baru | 2009 | p<0.01 | 2,019 | 408.4 | 5.1 | 11 | 492,942 |
| 3 | Colon | 2009 | p<0.01 | 1,402 | 188.2 | 7.4 | 1 | 206,553 |
| 4 | Calobre | 2015-2017 | p<0.01 | 2,120 | 511.5 | 4.1 | 4 | 162,315 |
| 5 | Arraijan | 2005-2006 | p<0.01 | 1,923 | 608 | 3.2 | 1 | 220,779 |
| **Adjusting for *Ae. albopictus* presence & absence** |  |  |  |  |  |  |  |  |
| 1 | Balboa | 2013-2015 | p<0.01 | 5,846 | 1,636.70 | 3.9 | 5 | 368,341 |
| 2 | Colon | 2009 | p<0.01 | 1,402 | 178.4 | 8 | 1 | 206,553 |
| 3 | Changuinola | 2005-2007 | p<0.01 | 1,734 | 296.5 | 6 | 2 | 114,445 |
| 4 | Santa Maria | 2015-2017 | p<0.01 | 2,013 | 445 | 4.6 | 3 | 139,778 |
| 5 | Arraijan | 2005-2006 | p<0.01 | 1,923 | 591.4 | 3.3 | 1 | 220,779 |
| **Adjusting for *Ae. aegypti* presence & absence** |  |  |  |  |  |  |  |  |
| 1 | Balboa | 2013-2015 | p<0.01 | 5,846 | 1,318.90 | 4.9 | 5 | 368,341 |
| 2 | Colobre | 2015-2017 | p<0.01 | 2,019 | 544.6 | 4 | 4 | 162,315 |
| 3 | Colon | 2009 | p<0.01 | 1,402 | 247.2 | 5.8 | 1 | 206,553 |
| 4 | Baru | 2009 | p<0.01 | 2,120 | 535.3 | 3.9 | 11 | 492,942 |
| 5 | Capira | 2014 | p<0.01 | 1,923 | 745.9 | 3.1 | 10 | 652,859 |

**Table 2.** Characteristics of each space-time model.

| Model | Total number of districts | RR 0-1 (# of districts) | RR > 1 (# of districts) | Highest RR | Most observed cases |
| --- | --- | --- | --- | --- | --- |
| No covariates | 20 | 9 | 11 | Bocas Del Toro (5.2) | San Miguelito (13,109) |
| Both *Aedes* species | 22 | 12 | 10 | Santiago (2.9) | San Miguelito (13,109) |
| Only *Ae. albopictus* | 12 | 4 | 8 | Bocas del Toro (6.2) | San Miguelito (13,109) |
| Only *Ae. aegypti* | 31 | 17 | 14 | San Miguelito (3.3) | San Miguelito (13,109) |

Furthermore, cluster 2 was found in different geographic locations for the *Aedes* (both) and *Ae. albopictus* models. This variation in duration of the clusters between the four models is a result of adjusting for the presence of *Aedes* during the 13-year study period. In other words, the start, end, and duration of the clusters is substantially affected by the presence of one or more *Aedes* species. The relative risk may be higher if *Aedes* was found in a district during the entire duration of a space-time cluster. During the 13 years of our study period combined with the 63 districts containing data (13 * 63 = 819), *Ae. aegypti* was present 690 times, *Ae. albopictus* was present 245 times, while both *Aedes* species were found in a district 224 times. As a result, the difference in species presence during the study period partly explains why the clusters for the *Ae. albopictus* model contained 19 less districts than the *Ae. aegypti* model, and 10 less districts that the model adjusting for both species.

The results of our GLM indicate that dengue PR can be predicted by the presence of *Ae. aegypti* alone, with all other covariates exhibiting insignificant statistical relationships to PR (*P* > 0.05), with covariate selection employed. Thus, controlling for all other factors, districts with a presence of solely *Ae. aegypti* exhibited an increase in adjusted PR of 1.0933 (*P* = 0.001). Additionally, the presence of *Ae. albopictus* did not predict the presence of *Ae. aegypti* (*P* > 0.05).

## Discussion

Our non-spatial statistical testing complements the space-time models, highlighting the likely role of *Ae. aegypti* in dengue transmission dynamics across Panama as well as the need to incorporate vector data into systematic dengue risk projections. In the model where *Ae. aegypti* presence and absence was accounted for, more than double the number of districts were contained in clusters than the model where *Ae. albopictus* presence and absence was accounted for. The *Ae. aegypti* model also contained the highest number of districts with a relative risk > 1, indicating more dengue cases than would be expected given baseline population levels. Findings are further supported by our determination that *Ae. aegypti* is the only predictor of dengue PR in the non-spatial model, which holds important implications for the understanding of dengue transmission dynamics in the changing landscape of vector ecology. As an invasive species that has systematically replaced *Ae. aegypti* throughout numerous regions in its endemic range [14], *Ae. albopictus* has been spreading throughout Panama for the previous 13 years [12,13]. Our results illustrate that it has not been a key driver of dengue prevalence throughout its time occurring in the country, but that more importantly, there is reason to believe that dengue rates may decrease as the species further proliferates, extirpating *Ae. aegypti* from its resident range within Panama. Globally, while *Ae. albopictus* has been implicated in several small outbreaks [35], the majority of dengue serotypes are thought to be transmitted by *Ae. aegypti,* due to its preference for both urbanized habitat [16,36] and human hosts [37,38].

Perhaps curious is the lack of association found with the other covariates, which included the presence and absence of *Ae. albopictus*, coexistence of both species, and the three socioeconomic variables. There have been a number of studies addressing the vector status and potential of *Ae. albopictus*. While it is biologically capable of transmitting dengue fever [6], outbreaks that can be directly attributed to this species are rare [35,39–41]. The lack of contribution of socioeconomic variables is also interesting, given socioeconomic conditions have been found to influence vector distribution [42–44]. However, no clear connection has been found between dengue risk and particular socioeconomic conditions [45], thus supporting our results. Overall, based on our findings, we suggest that vector surveillance results be incorporated into vector control planning. Specifically, focusing on regions where *Ae. aegypti* still maintains a stronghold may be an effective way of combating dengue outbreaks. Balboa, for example, was identified as a cluster in all four models and had a steady presence of *Ae. aegypti* throughout the sample period as well as increasing presence of *Ae. albopictus* since 2006. This district is relatively rural with approximately 2400 people spread across 400km^2^ area. It is possible that vector control efforts in Balboa are not as frequent or efficacious as in the more populated regions, yet this hypothesis would require field testing to confirm. Another district, San Miguelito in metropolitan Panama City, contained the most observed cases during our study period, despite being only 49.9km^2^. This district can be characterized by high density housing and residents of relatively low socioeconomic status. The staggering number of cases should be a cause for concern, yet its small geographic area may facilitate public health interventions such as vector control and community education. Overall, now that the identification of high risk districts at the national scale has been completed and informed by vector presence, the subsequent step of illustrating the comparative characteristics of each district relative to dengue transmission risk can be undertaken. Understanding what caused Balboa and San Miguelito to experience such high relative risk, for example, is the next task necessary for adjusting public health interventions to effectively address the needs and conditions of each district.

Despite the longevity of our data and thoroughness of the surveillance efforts, there are clear considerations and limitations of our work which we would like to see addressed in future studies. First, it is possible that the reported cases of dengue in certain districts are travel cases (seeking treatment in a district different than actual residence), therefore, adjusting for the presence of *Aedes* can shed light on the districts where an individual is more likely to get infected with dengue, not necessarily where all total cases were recorded. The lack of population data for more than one year across such a lengthy period is a considerable shortcoming of this work. While the frequency of a census in Panama is on par with much of Latin America, this greatly impacts our ability to determine accurate prevalence rates year to year. Since linear interpolation is often inaccurate for non-linear trends like population growth rate, we would like to see more frequent population assessments conducted in regions where dengue is an ongoing risk, and while we understand that resources may not easily allow for this, the role of national census efforts in public health is often under-appreciated. Second, the cylindrical shape of the clusters does not represent the true shape of the clusters, while it is possible to use irregular search windows [46–48]. Third, the STSS reports the relative risk for the entire study period, while relative risk will likely vary temporally. A final core limitation is the vector surveillance methods employed. Values are reported as the number of houses containing larvae of each respective species. No information is given on the number of houses surveyed, and thus we were forced to transform the data into presence and absence. Had the total number of surveyed houses been reported, we would have been able to compute each district’s infestation rate, which would have provided a scaled and more nuanced independent variable to compare to dengue PR.

Overall, it is key to recognize that adding vector surveillance data as a covariate changes the location, duration, and relative risk of dengue case clusters. Although unadjusted cluster analysis is a valuable tool for public health officials to identify high risk areas of vector-borne disease, our study illustrates the role that incorporating relevant covariates can play in altering the model output. While this has been demonstrated in cancer [49,50], this is the first use of covariates in space-time cluster detection modelling of neglected tropical disease. With this comes potential to expand into other classes of covariates. For example, in addition to vector surveillance data, we support the incorporation of additional covariates such as vector genetic background, climate, vegetation, and land cover to dengue cluster models. Furthermore, the differences reported for the clusters of dengue after adjusting for vector presence merit further small-area studies to determine local-scale characteristics that may assist in targeted intervention campaigns. Vector surveillance clearly provides valuable information in the determination of virus case clusters, and thus should be conducted alongside virus surveillance so that it may be included in modelling efforts. We intend for this exploratory study to inspire future investigations into the vector status of *Ae. albopictus* as well as the role of vector surveillance in public health planning efforts. We hope Panama’s robust dengue surveillance program can stand as a model for practitioners elsewhere, where current surveillance may be less thorough.

## Acknowledgements

We acknowledge Project Mosaic at the University of North Carolina at Charlotte for their assistance in data analysis. We also acknowledge Dr. Owen McMillan at the Smithsonian Tropical Research Institute (STRI), who was oversaw the project and its administrative responsibilities. We thank the staff of the Department of Statistics and Vector Control of MINSA for providing entomological data on *Aedes* mosquitoes as well as case counts for dengue fever across Panama over the study period. We are grateful to Panama’s Environmental Authority (*Mi Ambiente*, formerly ANAM) for supporting scientific collecting of mosquitoes.

## Funding

This work was partially supported by Secretaria Nacional de Ciencia, Tecnología e Innovación de Panamá (SENACYT-IDDS15_047) to JRL, http://www.senacyt.gob.pa/. JRL also further supported by SENACYT’s National Research Investigation System (SNI). The funders had no role in study design, data collection and analysis, decision to publish, or preparation of the manuscript.

